# SCALA: A web application for multimodal analysis of single cell next generation sequencing data

**DOI:** 10.1101/2022.11.24.517826

**Authors:** Christos Tzaferis, Evangelos Karatzas, Fotis A. Baltoumas, Georgios A. Pavlopoulos, George Kollias, Dimitris Konstantopoulos

**Author notes:** Georgios A. Pavlopoulos; George Kollias, Dimitris Konstantopoulos; Biomedical Sciences Research Center “Alexander Fleming”, 34 Fleming Street, Vari, 16672, Greece.

## Abstract

Analysis and interpretation of high-throughput transcriptional and chromatin accessibility data at single cell resolution are still open challenges in the biomedical field. In this article, we present SCALA, a bioinformatics tool for analysis and visualization of single cell RNA sequencing (scRNA-seq) and Assay for Transposase-Accessible Chromatin using sequencing (ATAC-seq) datasets. SCALA combines standard types of analysis by integrating multiple software packages varying from quality control to identification of distinct cell population and cell states. Additional analysis options enable functional enrichment, cellular trajectory inference, ligand-receptor analysis and regulatory network reconstruction. SCALA is fully parameterizable at every step of the analysis, presenting data in tabular format and produces publication-ready 2D and 3D visualizations including heatmaps, barcharts, scatter, violin and volcano plots. We demonstrate the functionality of SCALA through two use-cases related to TNF-driven arthritic mice, handling data from both scRNA-seq and scATAC-seq experiments. SCALA is mainly developed in R, Shiny and JavaScript and is available as a web application at http://scala.pavlopouloslab.info or https://scala.fleming.gr.

## INTRODUCTION

Single cell RNA sequencing (scRNA-seq) and ATAC sequencing (scATAC-seq) are both Next Generation Sequencing (NGS) techniques that have enabled the study of the transcriptome and epigenome respectively at an unprecedented resolution (1–5). Exploitation of these two modalities allows researchers to observe the heterogeneity of cell populations in more depth compared to established bulk RNA-sequencing techniques.

Since the first scRNA-seq publication (6), advances in technology and equipment have led to an exponential increase in the number of cells (from hundreds to millions) which can be simultaneously sequenced in one run. Widely used technologies which have been introduced over the past ten years include Fluidigm C1 (7), Smart-seq2 (8), Drop-seq (9) and 10x Genomics (10), whereas new protocols such as the 10x multiome and spatial transcriptomics (11) have also emerged. Both scRNA-seq and scATAC-seq techniques have been used in various experimental settings such as the investigation of different tissues, developmental timepoints, disease states and organisms. ScRNA-seq for example, has played a crucial role in the comprehensive annotation of cell types in multiple organisms (e.g., Human Cell Atlas for *Homo sapiens* (12), Tabula Muris for *Mus musculus* (13)), as well as in the identification of novel cell populations, sub-populations and disease states (e.g., (14)). Similarly, scATAC-seq has contributed critically in determining cell types in even higher resolution, as well as the epigenetic landscape that drives cellular differentiation, by characterizing gene regulation, and inferring Gene Regulatory Networks (GRNs) in several species and disease systems. Characteristic example of scATAC-seq analysis milestones is the case of *Drosophila melanogaster* brain epigenetic profiling (15), as well as the characterization of the chromatin accessibility profiles of 30 tissue types in Human (16).

From data generation to analysis and interpretation, a thorough bioinformatics pipeline is essential (14). Typical steps of such analyses include: *(i)* Quality Control (QC), *(ii)* read mapping and counting, *(iii)* normalization, *(iv)* dimensionality reduction, *(v)* clustering, *(vi)* differential expression/accessibility, *(vii)* peak calling, *(viii)* functional enrichment, *(ix)* data integration, *(x)* trajectory inference, *(xi)* generation of GRNs and *(xii)* visualization of results at every step. To this end, several software applications and packages which implement the aforementioned tasks have been proposed (17). Seurat (18), Scanpy (19), Monocle (20, 21), Cicero (22), Signac (23), EpiScanpy (24), SCENIC (25), cisTopic (26) and ArchR (27) are widely used R and Python libraries, whereas software which come with a graphical user interface (GUI), include Scope (28), Azimuth, Cerebro (29), iCellR (30) and SeuratWizard (31).

Seurat and Scanpy have been primarily used for the analysis of scRNA-seq data, offering functionalities varying from QC to population identification and integration of multiple datasets, while Signac and EpiScanpy extend Seurat’s and Scanpy’s functionality to process scATAC-seq data. ArchR focuses on the analysis of single cell chromatin accessibility data by offering standard analysis steps, as well as additional powerful features like Positive Regulator identification, Transcription Factor (TF) footprinting and trajectory inference. Monocle is a scRNA-seq analysis package that offers a widely used pseudo-temporal cell ordering framework, while its extension, Cicero can be used for the analysis of scATAC-seq data. Regarding tools with a GUI, Scope offers various visualization options including a side-by-side comparative view at cluster and gene levels for datasets containing multiple samples, disease conditions or timepoints. While this is practical for exploring clustering results, GRNs and expression patterns, the tool lacks further downstream data analysis. Azimuth focuses on the basic scRNA-seq analysis steps, but lacks customization options as it mainly specializes in the characterization of the identified populations by adopting a ‘reference-based mapping’ approach. SeuratWizard exploits the standard steps of the analysis, while Cerebro also builds upon the initial results, allowing the user to explore additional modes such as signature scoring, cell cycle phase analysis and trajectory inference. Finally, iCellR covers both scRNA-seq and scATAC-seq basic analyses but lacks ligand-receptor and GRN reconstruction.

In this article we present SCALA, a holistic pipeline which integrates all the aforementioned procedures and enables biomedical researchers to get actively involved in the downstream analysis and exploration of both scRNA-seq and scATAC-seq datasets. SCALA is a fully interactive bioinformatics tool which offers access to all standard analysis modes, varying from QC and data normalization to the identification of distinct cell populations and cell states. Furthermore, SCALA supports additional analysis modes such as automatic cluster annotation, functional enrichment analysis, ligand-receptor analysis, trajectory inference and reconstruction of GRNs. Interactive plots as well as publication-ready figures and data tables can be generated at every step of the analysis while any of the processed datasets can be exported to be further analyzed with external applications. We believe that SCALA can become a go-to tool for experimentalists who seek to analyze their scRNA-seq and scATAC-seq datasets and communicate biological findings with high resolution visualizations.

## METHODS

### Input data types

SCALA is compatible with several input data types. For scRNA-seq, the primary data input consists of a unique molecular identifier (UMI) count matrix. The user can provide such a matrix by either uploading a gene (rows: features) by cell (columns: barcodes) tab-delimited data table (including row and column names), or by uploading the output of the cellranger pipeline from *10X* (filtered_bc_matrix). In the latter case, the cellranger count output folder should contain: *(i)* a file named “barcodes.tsv.gz” containing only detected (filtered by cellranger count pipeline) cellular barcodes in gzip CSV format, *(ii)* a file named “features.tsv.gz” with features (genes) that correspond to row indices in gzip TSV format; the columns of the particular file should correspond to feature ID, feature name and feature type (Gene expression) respectively, *(iii)* a feature-barcode count matrix in gzip Market Exchange Format (MEX). Moreover, the user has a third option of uploading a pre-analyzed *Seurat* object in RDS (*R* saved object) format. In the case of scATAC-seq, SCALA is only compatible with arrow files in its current version. The particular file format stores all the associated data (i.e., metadata, accessible fragments and data matrices) within a sample. Arrow files can be created by using the *create_arrow_file*.*R* helper script provided in SCALA’s GitHub repository or by using the *ArchR* package directly.

### Functionality

After the input files have been uploaded, SCALA’s main workflow (Figure 1) can be utilized for both scNGS pipelines. The steps are: *(i)* QC, *(ii)* data normalization and scaling, *(iii)* variable features detection, *(iv)* Principal Component Analysis (PCA) dimensionality reduction, *(v)* Latent Semantic Indexing (LSI) dimensionality reduction, *(vi)* clustering, *(vii)* additional dimensionality reduction methods, *(viii)* feature inspection, *(ix)* markers’ identification, *(x)* cell cycle phase analysis, *(xi)* functional/motif enrichment analysis, *(xii)* clusters’ annotation, *(xiii)* trajectory analysis, *(xiv)* Ligand-Receptor (L-R) analysis, *(xv)* GRN analysis and *(xvi)* visualization of epigenome signal tracks.

**Figure 1:**
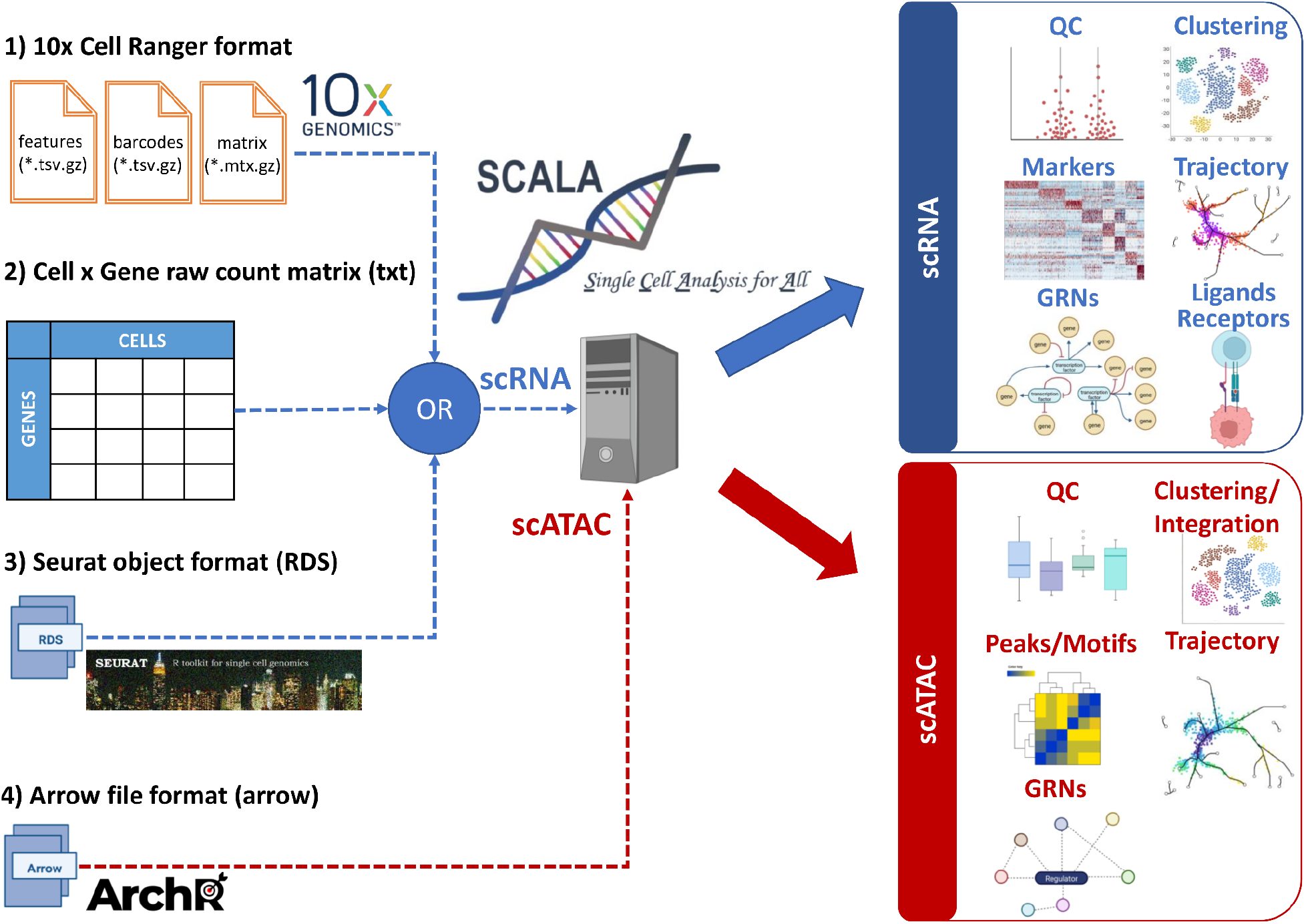
General workflow of the SCALA pipeline. In this figure the input files compatible with SCALA for scRNA-seq and scATAC-seq analysis are shown in the left panel. Additionally, the main functionalities and outputs, for each mode of analysis for RNA (blue box) and ATAC (red box) assays, are showcased in the right panel.

### Quality Control

Identification and removal of “low quality” cells (empty, stressed, broken, or dead cells) and non-informative genes is essential for downstream analysis in scNGS datasets. SCALA allows the exploration of QC plots and filters-out cell barcodes through application of user-defined parameter thresholds. Common scRNA-seq QC criteria include *(i)* per cell number of unique features detected, *(ii)* per cell detected UMIs and *(iii)* per cell percentage of mitochondrial content. Cells that exhibit very low numbers of *(i)* and *(ii)* are typically excluded as low-quality, while those with very high numbers are considered as putative multiplets. Barcodes with a high percentage of mitochondrial UMIs should also be excluded as low-quality/dying cells (Figure 2A).

**Figure 2:**
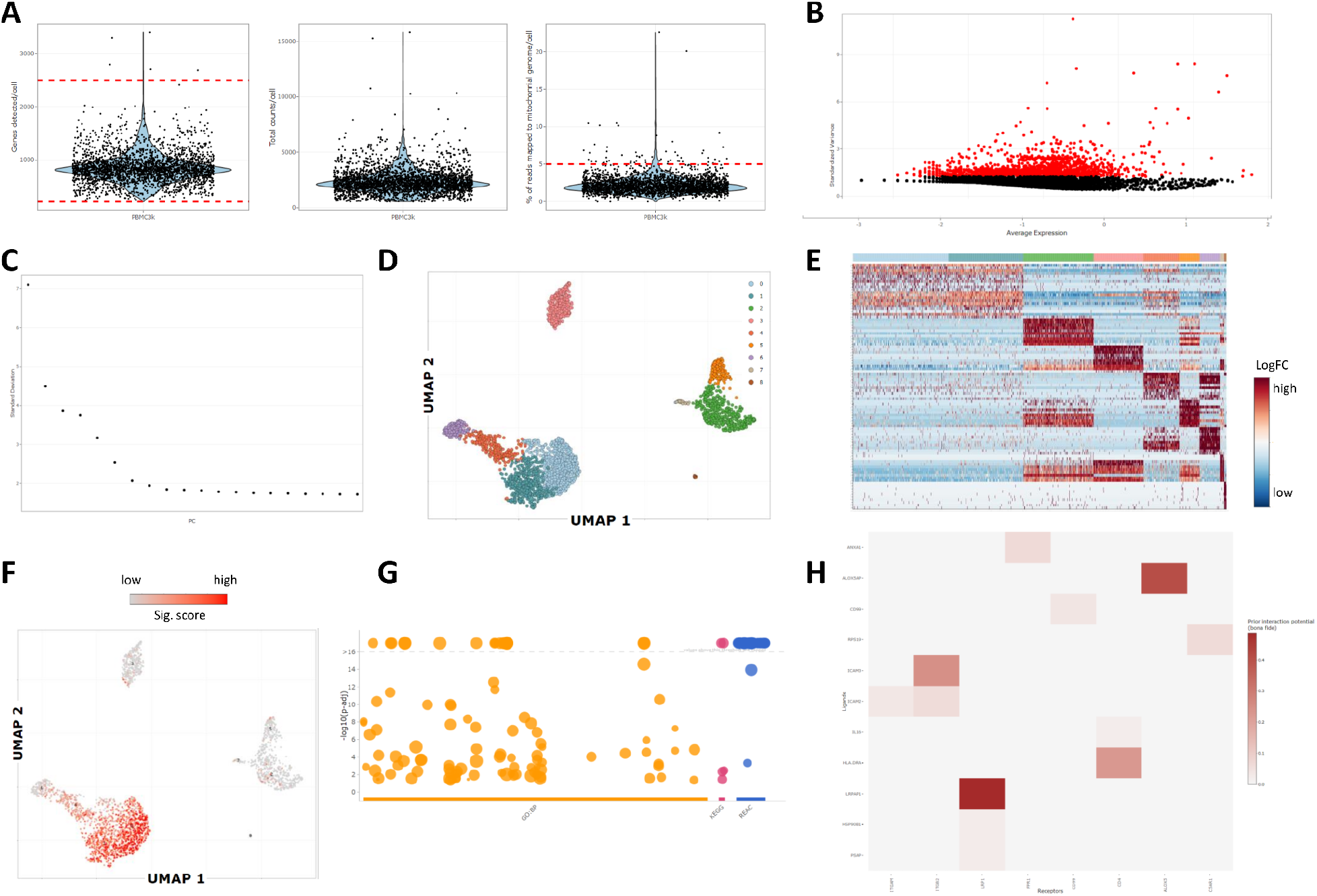
Analysis of PBMC3k scRNA-seq dataset. (A) Violin plots depicting cell quality control measurements including the number of genes detected, the total number of reads and the percentage of reads mapped to the mitochondrial genome. (B) Detection of most highly variable genes using the VST selection method. (C) Scatterplot for the top 20 Principal Components computed by PCA. (D) Visualization of cells in UMAP space. Cells are colored according to cluster labels (clusters were identified with the Louvain algorithm). (E) Heatmap depicting the top10 marker genes per cluster, ranked by Log2FC value. (F) Feature plot showcasing signature scores/per cell for the top marker genes of cluster 0. (G) Manhattan plot depicting enriched GO:BP, KEGG and Reactome pathways for the positive marker genes of cluster 0. (H) Heatmap showing “bona fide” interactions between clusters 0 (ligand expressing cluster) and 2 (receptor expressing cluster). The intensity of the color represents the interaction potential score.

Similarly, typical scATAC-seq QC metrics include: *(i)* transcription start site (TSS) enrichment and *(ii)* the number of unique nuclear fragments (*log10(nFrags)*). TSS represents the chromatin accessibility signal-to-background ratio. Enrichment of ATAC-seq signal in TSS regions of expressed genes is typically high in most cell types and a classic criterion of the quality of the assay. The particular metric is calculated as the ratio between TSS enrichment relative to per-base pair 2 kb flanking regions enrichment. Furthermore, cells including too few nuclear fragments should be excluded in order to avoid inclusion of non-interpretable data (Figure 3A).

**Figure 3:**
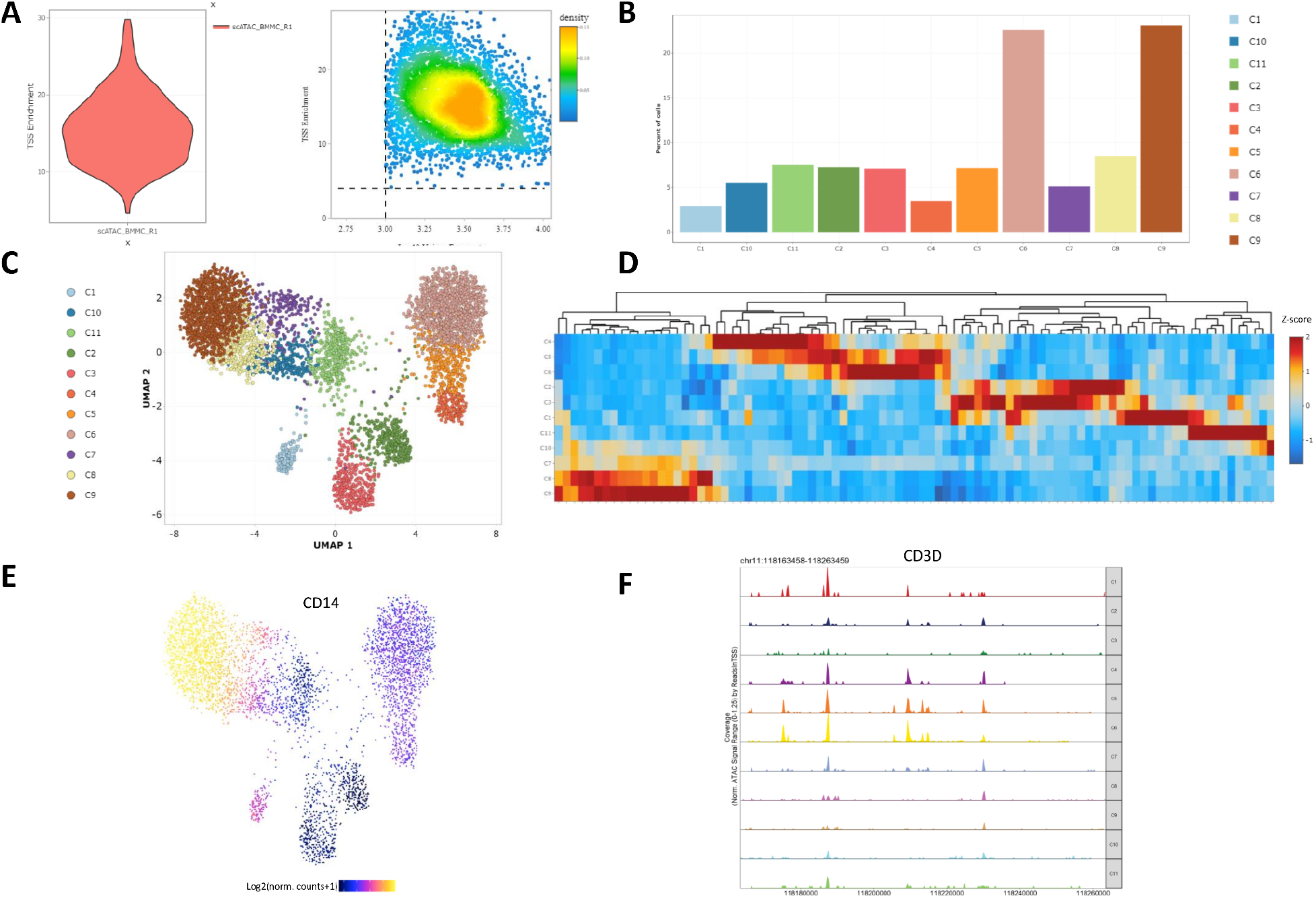
Analysis of BMMCs scATAC-seq dataset. (A) Cell quality control plots depicting information about TSS enrichment and unique fragments measurements. (B) Bar plot showing the relative abundance of cells in each of the dataset’s clusters (clusters were identified with the Louvain algorithm). (C) Projection of cells in UMAP space. Cells are colored according to cluster identity. (D) Heatmap showing z-scores of peak accessibility for the top marker peaks per cluster. (E) Feature plot showcasing gene activity scores (per cell) of CD14 as a UMAP overlay. (F) Genome browser tracks showing local chromatin accessibility of CD3D gene at cluster level.

### Data normalization and scaling

scRNA matrices are normalized and scaled in order to eliminate cell-depth variability biases as well as to transform the data properly prior to variable feature detection and dimensionality reduction. Data normalization in SCALA is applied through a global-scaling normalization method (18), where the gene count of each barcode is normalized by the total barcode counts, multiplied by 10,000 and log-transformed. Normalized values are stored in a *Seurat* object, and normalized counts are further standardized to z-scores, with column-wise mean expression equal to 0 and variance equal to 1. In order to mitigate the effect of unwanted sources of variation, the user can optionally provide metadata variables. In such a case, they are individually regressed against each feature, while scaling and centering is then performed on the resulting residuals.

### Variable features detection

Using the normalized RNA data matrix, genes that exhibit the highest column-wise variation are detected. Targeted analysis of the particular subset of features aids in the identification of the underlying biological patterns in single cell datasets. The supported methods for the identification of most variable features (Figure 2B) include three methods. These are *(i)* Variance Stabilizing Transformation (VST), *(ii)* Mean-Variance Plot selection (MVP) and *(iii)* “Dispersion”. VST fits a line in the relationship of log-variance and log-mean using local polynomial regression. Consequently, standardization of feature values using the observed mean and expected variance is performed. Feature variance is finally calculated for standardized values, after clipping to a maximum. A fixed number (default = 2,000) of variable features is returned. MVP uses a function to calculate average gene counts and gene dispersions. In this function, all genes are separated into 20 bins according to their average counts. Finally, dispersion z-scores are calculated for each gene group. For “Dispersion”, genes with the highest dispersion values are kept. For the last two methods, a variable number of features is returned.

### PCA dimensionality reduction

PCA is performed on the scaled values of the most variable features to determine the “dimensionality” of the dataset. The most informative Principal Components (PCs) are identified and used in the next steps of cell clustering and cluster visualization. The number of PCs that exhibit the higher variation of the scRNA matrix can be determined either automatically by applying *10-fold Singular Value Decomposition* (SVD) cross validation or customly by examining the ranking of the incremental variance of each PC (elbow plot, Figure 2C).

### LSI dimensionality reduction

LSI is performed in scATAC-seq matrices, using genome-wide 500-bp tile counts (27). Tile-counts are normalized to eliminate the cell depth bias using a constant of 10,000, followed by inverse document frequency normalization and log-transformation. During this process, the most variable features (tiles) are detected. The particular process is run in an iterative manner where an LSI transformation is applied using the most accessible features (tiles). This procedure identifies lower resolution clusters that are not batch confounded. Consequently, average accessibility for each of these clusters is calculated across all features. Finally, the most variable features are identified across low resolution clusters and are used as input for the next LSI iteration.

### Clustering

Graph-based clustering is performed in scRNA-seq (Figure 2D) and scATAC-seq (Figure 3B) matrices, in order to define cell types and/or cellular states. Initially, cells are embedded in a *Shared-Nearest Neighbor* (SNN) graph structure based on Euclidean distances in the PCA/LSI space. Cells that exhibit similar gene expression/chromatin accessibility profiles are connected with edges. The newly formed graph is then partitioned into highly interconnected communities using the Louvain algorithm (32).

### Additional dimensionality reduction methods

To visualize cells, cell clusters and cluster relationships in 2D and 3D space, additional dimensionality reduction techniques, are applied, like uniform manifold approximation and projection (UMAP) (33) (Figure 3C), t-Distributed Stochastic Neighbor Embedding (tSNE) (34), diffusion maps (35) or Potential of Heat-diffusion for Affinity-based Trajectory Embedding (PHATE) (36). Such visualizations uncover the underlying modularity of the datasets. Additionally, they can be utilized for feature inspection, exploration of cluster structures and for trajectory inference purposes (especially PHATE).

### Markers’ identification

Differential expression as well as differential accessibility analysis enable the identification of marker genes (Figure 2E) and peaks (Figure 3D) respectively and guide the cell-type and cell-state annotation/characterization of cell clusters. Differential analysis assists the detection of key transcriptional and regulatory programs that drive pathogenicity and/or development. The analysis is performed in a cluster-specific manner, where each cluster’s cells are tested against all the other cells of the dataset. The available statistical tests for scRNA-seq are: *(i)* Wilcoxon rank sum test, *(ii)* likelihood-ratio test for single cell feature expression (37), *(iii)* standard AUC classifier, *(iv)* Student’s t-test, *(v)* MAST (38) and *(vi)* DESeq2 (39). Similarly, for scATAC-seq the tests are: *(i)* Wilcoxon rank sum test, *(ii)* Student’s t-test and *(iii)* binomial test.

### Feature inspection

Feature expression and chromatin activity can be explored by cell scatter plots (Figure 2F, Figure 3E) in reduced space (e.g., UMAP, tSNE, etc.), or via violin plots, heatmaps and dotplots. In scRNA-seq datasets, gene signatures can also be calculated by utilizing the UCell package and visualized as described above. Moreover, QC metrics such as total number of reads per cell, genes detected per cell, etc. can be visualized via scatter plots and violin plots at a cluster level.

### Cell cycle phase analysis

Calculation of cell cycle phase scores is based on S, G2/M and G1 canonical markers in the scRNA-seq count matrix. In cases where cluster-specific patterns of cell cycle biases are captured, the user has the option to use the “regress out” option (in the step of scaling) in order to mitigate the cell-cycle effect. The results of the analysis can be viewed either in a scatter plot format, where cells are projected in a reduced space (PCA, UMAP, tSNE, diffusion map, PHATE) and colored according to the predicted phase of cell cycle, or as a bar plot summarizing the percentages of cells assigned to each cell cycle phase per cluster.

### Functional/Motif enrichment analysis

Using the previously identified marker genes and peaks, functional enrichment analysis (e.g., for pathways and Gene Ontologies (GOs)) and motif enrichment analysis can be performed for each cluster. In detail, for scRNA-seq data, up/down regulated genes from the clusters identified in previous steps are tested for enriched GO terms or KEGG pathways, using the g:Profiler package (40). The enriched terms can be visualized in a table format accompanied by information about statistical significance and gene overlap (between the input list and the term of interest). Additionally, a bubble plot summarizing the enriched terms per database used is also available (Figure 2G). Regarding motif enrichment analysis, marker peaks identified in previous steps are tested for enrichment of binding sites of specific transcription factors (TFs). Finally, deeper functional enrichment analysis with more informative visualization is also offered by Flame (41) external application. This can be done per cluster (one gene list), or multiple gene lists (up to 10 clusters) can be analyzed simultaneously with the help of interactive UpSet plots.

### Cluster annotation

For automated cluster annotation, the CIPR package is utilized (42) as it contains reference datasets for human and mouse organisms, to assign cell-type identities. The end user can select a dataset to use as a reference and the type of analysis that will be employed to calculate the predictions, either by keeping all dataset genes or only the differentially expressed ones. Moreover, the user can also select the correlation metric (Pearson, Spearman) that will be used. Regarding the visualization options, a table containing all predictions per cluster is returned, as well as a dot plot that summarizes the top-5 predictions per cluster.

### Multimodal Integration analysis

In this mode of analysis, the user can upload an already processed scRNA-seq dataset to perform integration analysis with the scATAC-seq dataset, which is currently loaded in SCALA. More specifically, gene activity scores from the ATAC assay and gene expression values from the RNA assay are combined in order to align cells between datasets. The output of integration analysis results in transferring labels from scRNA clusters to cells from the scATAC dataset. The newly obtained clustering identities of the cells can be adopted in other downstream steps such as detection of marker peaks, trajectory analysis, etc.

### Trajectory analysis

Pseudotemporal ordering of single cells, facilitates the uncovering of underlying differentiation/developmental processes which lead cells to transitions between different cellular states. In SCALA, Slingshot (43) is employed, utilizing input clustering information and dimensionality reduction coordinates for all cells of a dataset, in order to construct a Minimal Spanning Tree (MST) at the cluster level. The nodes of the tree represent the clusters while the edges represent their in-between relationships. The user can select the dimensionality reduction method which will be used for the Slingshot execution (PCA, UMAP, tSNE, diffusion map or PHATE), as well as the initial and final states, which define the direction of the identified trajectory. The MST is always drawn in a UMAP plot, while pseudotime values are calculated per lineage and can be visualized again in UMAP space as a separate scatter plot.

### Ligand - Receptor analysis

The prediction of ligand-receptor interactions is a crucial step for deciphering cell-to-cell communication at different tissues. Inspection of communication patterns between different cell types could aid in the detection of key interactions, driving gene expression alterations (downstream of signaling pathways) at healthy and disease contexts. SCALA, incorporates the analysis framework of nichenetR (44). More specifically, after clustering the user needs to select a pair of clusters that will be used to search for L-R interactions. As a first step, overexpressed genes are calculated in each cluster. Then the reported interactions are ranked, by considering a “prior interaction potential” score that is calculated in the initial steps, when the protein-protein interaction model is constructed. A heatmap visualization summarizes all the interactions that have been detected between the two clusters of interest (Figure 2H). L-R interactions and their respective scores are available for download in a table format including the “prior interaction potential” score that signifies the strength of the predicted interaction.

### Gene Regulatory Network analysis

In this step, by utilizing the SCENIC workflow (25), co-expression modules of TFs (Transcription Factors) and their target genes are detected based on co-expression analysis and TF motif analysis. Area Under the Curve (AUC) scores per cell are calculated and denote the activity of a regulon, defined as a group of genes containing a TF and its target genes. Finally, average AUC values and Regulon Specificity Score (RSS) scores, which showcase the activity and specificity of regulons, can help the user visualize active regulatory networks in heatmap format and examine whether cluster specific regulons are present in the dataset. Due to limitations in run-time in *R* environments, SCALA offers instructions so that an end-user can externally run some parts of the analysis in *Python* and then import the result files again in SCALA for visualization. Gene regulation analysis at chromatin level, aims to identify cluster specific TFs, whose expression exhibit a high correlation with chromatin accessibility changes at genomic sites, that include their DNA binding motifs (positive regulators).

### Visualization of epigenome signal tracks

Chromatin accessibility tracks can be used as an alternative to feature plots (which depict gene scores in reduced space). The user can select a gene and the number of bases upstream and downstream, defining a genomic interval of interest. The inspection of the plot (Figure 3F) through a genome browser snapshot can reveal chromatin accessibility in the gene body or upstream/downstream gene regulatory elements (promoters, enhancers, silencers, etc.).

### Implementation

SCALA is mainly developed in R/Shiny and JavaScript. For the basic analysis of scRNA-seq data, the Seurat package is utilized, while ArchR is employed for the basic steps of scATAC-seq analysis. Furthermore, downstream applications including functional enrichment analysis, trajectory inference, GRNs reconstruction and L-R analysis were made possible by the incorporation of the g:Profiler (40), Slingshot (43), SCENIC (25) and nichenetR (44) packages. In the online version, the supported dataset size is restricted to <2GB while no more than 2 CPU threads on the server are allocated per session, something that may result in slow execution. However, by downloading the standalone version of SCALA from GitHub, users can bypass both aforementioned restrictions (settings variables in global.R file).

## RESULTS

### Analysis of two datasets for synovial fibroblasts in arthritis mouse model

To demonstrate SCALA’s functionality, we utilized two previously published datasets (45) (scRNA-seq and scATAC-seq) regarding the single cell transcriptome and chromatin dynamics of Synovial Fibroblasts transitioning from homeostasis to pathology in modeled TNF-driven arthritis. For this purpose, the human TNF (*hTNFtg*) transgene overexpressing mouse model Tg197 (46) was used and compared against healthy wild type (Wt) mice. As reported previously (45), for the generation of *10x Genomics* scRNA-seq libraries (Single Cell 3’ v3 reagent kits), 6,667 sorted non-hematopoietic stromal cells (*Cd45*-, *Cd31*-, *Ter119*-, *Pdpn*+) were isolated from whole ankle joint synovium. These libraries were sequenced at a depth of 400 million reads, using one lane of an Illumina NextSeq 500 machine. For the second dataset, scATAC-seq libraries were generated using a similar experimental set-up, according to *10x Genomic*s guidelines, profiling a total of 6,679 single nuclei. In each experiment, cells were derived from three healthy mice tissues (WT, 4 weeks of age), and six diseased *hTNFtg* mice; three at an early disease stage (*hTNFtg*/4, 4 weeks of age) and three at an established pathological stage (*hTNFtg*/8, 8 weeks of age). As shown in previous publications (46, 47), Tg197 mouse model over-expresses human TNF (huTNF) transgene, leading to the development of an arthritic phenotype manifested by cartilage destruction and bone erosion, which ultimately results in loss of joint function. Here, SCALA was used to reanalyze 5,903 synovial fibroblast (SFs) transcriptomes and 6,046 epigenomes, originating from healthy mice (control sample) and arthritic mice at 4 and 8 weeks of age (early and established disease state).

### Analysis using SCALA’s scRNA-seq pipeline

For the scRNA-seq QC step, cells with <500 features (genes) detected or having >10% of their reads mapped to the mitochondrial genome, were excluded from further analysis. Consequently, downstream analysis of scRNA-seq was performed as follows: Most highly variable feature detection was performed by applying the mean-variance-plot (MVP) method implemented by the Seurat package, leading to the identification of 1,535 variable genes. Gene counts of each cell were divided by the total cell counts, multiplied by 10,000, and natural-log transformed. Scaling of the normalized expression values was performed on all genes by utilizing the option of “regressing out” the mitochondrial reads effect.

The scaled gene-by-cell expression matrix of most variable genes was used as input to perform Principal Component Analysis (PCA). To identify the dimensionality of the dataset (most informative principal components in terms of cell heterogeneity), Singular Value Decomposition (SVD) *k*-fold cross-validation was performed using the dismo R library. This procedure determined the number of most informative principal components (25 PCs), which were used for the steps of cell clustering and non-linear dimensionality reduction analysis. Specifically, in order to identify distinct fibroblast subsets, graph-based clustering analysis was performed with Seurat’s Louvain algorithm, by setting the resolution parameter to 0.6. The 25 most informative PCs were also used for non-linear dimensionality reduction analysis (tSNE and UMAP), to visualize the newly formed cell clusters in 2D/3D space.

SF clustering led to the formation of 10 SF clusters, with distinct transcriptional profiles, exhibiting homeostatic, inflammatory and destructive characteristics in healthy and arthritic joints respectively. These characteristics were detected by performing marker gene identification analysis for each identified SF cluster. More specifically, each cluster’s transcriptomes were compared against the rest of the cells’ transcriptomes, through the Wilcoxon rank sum test on the normalized expression values. Genes with average log Fold Change (logFC) > 0.25, a percentage of expression/cell-occurrence (gene detected in a cell) > 25%, and a p-value < 0.01 were retained.

Consequently, up-regulated genes were used as an input to perform functional enrichment analysis. In particular, GO biological processes enrichment was conducted for each SF cluster, using g:Profiler (40). Examination of similarities/differences between SF clusters at the level of markers and enriched terms, led to the merging of two clusters, (0 and 9). The resulting nine clusters were designated as S1, S2a, S2b, S2c, S2d, S3, S4a, S4b and S5 (Figure 4A). It should also be pointed out that the identified clusters exhibit differences in their relative abundances in healthy and disease states. More specifically, a group of them is shrinking, while another is expanding during disease (Figure 4B). Thy1+ clusters (S1, S2a, S2b, S2c, S3 and S5) were further annotated as “sublining”. Interestingly, their transcriptional and functional characterization comprises features of tissue homeostasis preservation, with the exception of S5 which shows an immuno-regulatory role under healthy conditions. Enriched GO terms for these populations include BPM, WNT, TGFbeta and SMAD signaling pathways, as well as response to TNF and IFN-beta/gamma. Top markers for these clusters contain *Smoc2, Thbs1, Vwa, Rgma, Dkk2, Sfrp1, Ecrg4, Osr1, Nr2f2, Klf5, Clu, Id1, Meox1, Pi16, Sema3c, Efemp1, Ccl7, Il6*, and *Notch3*. Similarly, Prg4^High^ S4a cluster was annotated as “lining” and was linked with functions that define an inflammatory-destructive profile for the particular SF subpopulation. The lining phenotype is described by markers like *Tspan15, Hbegf, Htra4* and *Clic5*. Regarding the enriched biological processes we detected terms such as inflammatory response and class I antigen presentation. Finally, clusters S2d and S4b showed a mixed expression profile of *Prg4* and *Thy1* (Prg4+ Thy1+), and thus were annotated as “intermediate” subpopulations. Marker genes such as *Fbln7, Thbs4, Cthrc1, Lrrc15, Dkk3, Mki67, Pdgfa, Birc5, Aqp1, Acta2, Cxcl5*, which were found upregulated mainly in the intermediate and lining compartments, are either previously reported as players of fibroblast pathogenicity or linked to potential pathogenic roles. Corresponding terms like regulation of immune, redox response, fibroblast proliferation, cell division and apoptosis were found enriched in S2d and S4b. Conclusively, this group of clusters showcases a pro-inflammatory and proliferating character (Figure 4C).

**Figure 4:**
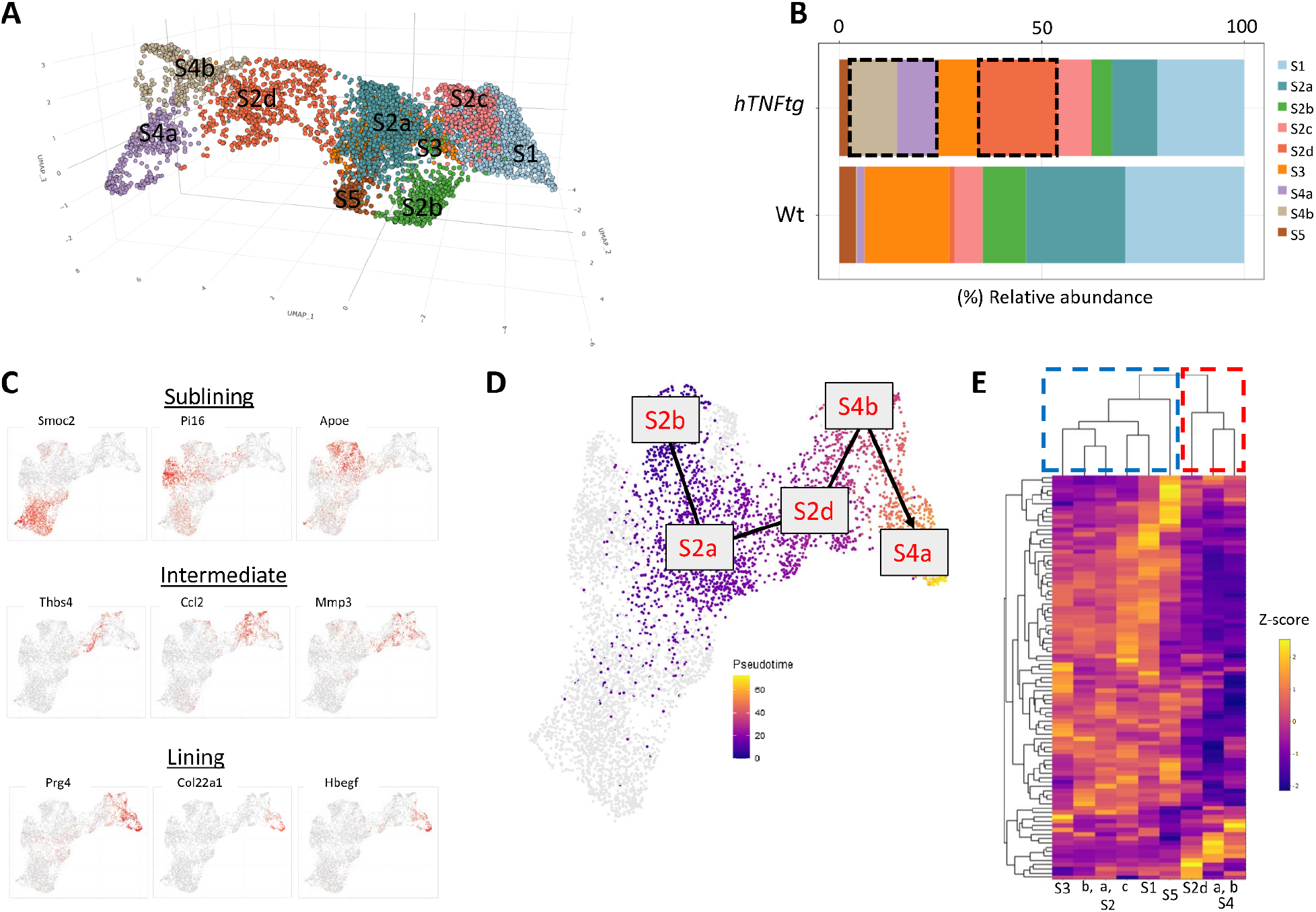
Use case - *hTNFtg* scRNA-seq data analysis. (A) Graph based clustering of SFs identified 9 distinct clusters. Cells are visualized in 3D UMAP space and are colored by cluster assignment. (B) Barplot depicts relative abundances of clusters in healthy (Wt) and disease (*hTNFtg*) states. The highlighted areas pinpoint the clusters that are expanded in arthritic state. (C) Feature plots showing the different gene expression patterns between the clusters of sublining (top row), intermediate (middle row) and lining (bottom row) categories. Cells are projected in the 2D UMAP space and colored by normalized gene expression. (D) One of the possible lineages (proposed by trajectory analysis) is showcased in UMAP overlay. Cells belonging to the lineage are colored according to their pseudo-time values, while cells that are not part of this lineage are colored in light gray. (E) Heatmap depicting regulon activity of top-80 regulons (z-scores of AUC values) at cluster level. Hierarchical clustering of fibroblast subsets (using active regulons) identified two major groups (group1: sublining clusters, group2: intermediate and lining clusters)

Next, cell cycle phase analysis was performed, assigning each cell in S, G1 or G2/M phase. Interestingly, one of the three SF populations exhibiting pathogenic characteristics (S4b), showed the highest percentage of cells located to G2/M phase. This finding was also supported by cycling markers (extracted from the literature), that were specifically expressed in the S4b cluster. The mixed expression signature of *Prg4* and *Thy1* (Prg4+ Thy1+), which characterize this “intermediate” group of cells, is thus a strong marker of disease state that is observed mainly in *hTNFtg* conditions.

Cellular trajectories were calculated for the pooled dataset using as an input to the slingshot algorithm the first 25 most informative PCs. In order to determine the clusters used as an input for the initial and final state of the trajectory, current literature (48, 49) was taken into consideration as well as the results of external software applications such as scVelo (50) and CellRank (51). The produced minimum spanning tree highlighted the existence of a pathogenic branch composed by clusters S2a, S2d, S4b, S4a, indicating S4a as final state and S1, S2b, S3 and S5 as potential starting points (Figure 4D).

We next sought to study ligand-receptor interactions between the sublining and intermediate compartments with lining. By employing the nichnetR package and focusing on ligands/receptors with a percentage of expression >10% in the clusters of interest, we identified shared and specific interactions. More particularly, we detected 157 and 152 interactions between sublining-lining and intermediate-lining respectively. In more detail, 126 of those interactions were shared, however 26 were specific to intermediate-lining and 31 to sublining-lining. Interestingly, in the interactions of sublining and lining we noticed pairs of ligands and receptors participating in Wnt and BMP signaling. In contrast, in the intermediate-lining interactions, we have detected pairs related to MMP13, IL-11 and RSPO2 signaling.

As a last step in the scRNA-seq analysis pipeline, GRN analysis was performed to detect regulons that exhibit preferential activation patterns at cluster level. That resulted in the identification of 133 regulons in total. Interestingly, distinct activation patterns were observable in the different clusters and hierarchical clustering of the top-80 regulons revealed two groups, the first containing only sublining clusters and the second containing intermediate and lining (Figure 4E).

### Analysis using SCALA’s scATAC-seq pipeline

Regarding the scATAC-seq data, QC was initially performed and cells with Transcription Start Site (TSS) enrichment score <4 and count-depth <1,000 unique nuclear fragments were removed from downstream analysis. Next, LSI was employed using a resolution of 0.6, a total of 30 dimensions, 4 iterations and otherwise default settings. Additionally, UMAP projection was produced for the visualization of cells in 2D space. Gene activity scores were computed as the summed local accessibility of promoter-associated count-tiles in the proximity of each gene, adopting a distance-weighted accessibility model (27). In detail, count-tiles in the range of 100,000 bp of a gene promoter were aggregated using the following distance weight formula: 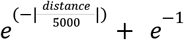. An extra normalization step was applied (multiplication by, 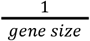, scaled linearly from 1 to 5), in order to account for gene length biases. As a following step, the above-weighted sum was multiplied by the aggregated Tn5 insertions in each tile. Gene scores were then scaled to 10,000 counts and log2-transformed. To improve the visualization of gene activity scores, a smoothing procedure was applied using the MAGIC algorithm (52).

Similar to the RNA analysis, clustering was performed with the use of the Louvain algorithm with resolution 0.6. This procedure led to the identification of 8 clusters (Figure 5A).

**Figure 5:**
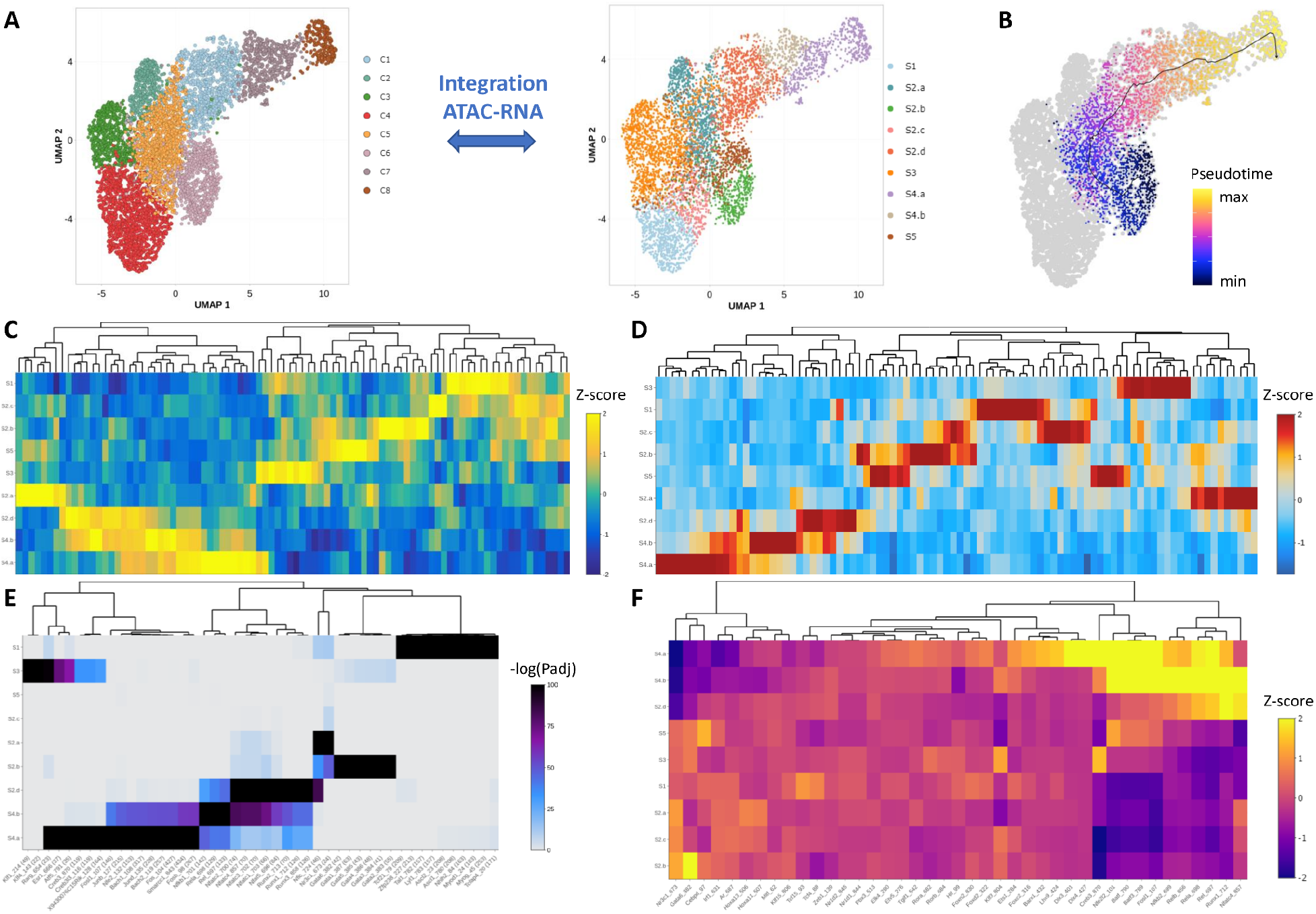
Use case - *hTNFtg* scATAC-seq data analysis. (A) Integration between scRNA-seq and scATAC-seq datasets. Cluster labels from RNA analysis are transferred to ATAC. Cells are projected in UMAP space and colored according to clustering (left) or transferred labels (right). (B) Semi-supervised trajectory analysis in ATAC dataset recapitulates the outcome of the respective analysis in RNA data. S2b was used as an initial state and S4a as a final state. (C) z-scores of gene activity for the top-10 marker features of each cluster (after integration) is displayed in a heatmap. (D) Heatmap displaying z-scores for the accessibility of top-10 marker peaks for each cluster (after integration). (E) Motif enrichment analysis in marker peaks of each cluster. Enriched motifs of each cluster are displayed in a heatmap. Color scale denotes the significance of enrichment. (F) Gene regulation analysis identifies positive regulators for each cluster. Top regulators are displayed in a heatmap.

Afterwards, integration between the ATAC dataset and the previously analyzed RNA dataset was performed. Our goal was to achieve “label transferring” between the annotated RNA clusters and the new groups that emerged after the ATAC clustering analysis. The integration process enabled the labeling of scATAC-seq cells according to the 9 SF subpopulations occurring in RNA analysis (Figure 5A) (differences in the software versions of Seurat and ArchR employed in SCALA, compared to the ones used in the publication containing the initial analysis of the dataset didn’t let us reproduce our UMAP visualization in the exact same manner).

Following integration analysis, semi-supervised trajectory inference with ArchR (Figure 5B), confirmed the existence of a pathogenic branch, consisting of S2a, S2d, S4b and S4a clusters, in accordance with scRNA findings.

By utilizing the gene activity scores (calculated as described above), Wilcoxon test was employed (Figure 5C) in order to detect top marker features per cluster (|Log2FC| ≥ 0.58 and FDR ≤ 0.05 cut-offs were applied). Consequently, a robust merged peak set was identified across SF clusters, using MACS2 (53) by generating two pseudo-bulk replicates. Iterative overlap peak merging (54) was applied at the level of the pseudo-bulk replicates and across SF subpopulations, forming a single merged peak set of 158,713 regions with a fixed length of 500 bps. Subsequently, differential accessibility analysis between cells was performed to identify cluster-specific marker peaks (|Log2FC| ≥ 0.58 and FDR ≤ 0.1 cut-offs were applied) (Figure 5D). Consequently, marker peaks were utilized to perform motif enrichment analysis, using the CIS-BP database (|Log2FC| ≥ 0.58 and FDR ≤ 0.05 cut-offs were applied) (Figure 5E). The three previous modes of analysis pinpointed the existence of distinct patterns of gene activity and peak/motif accessibility along clusters. Furthermore, hierarchical clustering on z-scores strengthened the categorization of clusters in three main groups namely sublining, intermediate and lining.

Gene regulatory analysis was conducted in the ATAC assay as well. More specifically peak to gene linkages were detected using correlation analysis between enhancer peak accessibility and integrated gene expression. Moreover, TF motif accessibility was correlated with integrated TF gene expression in a cell-by-cell manner, reporting TFs with Pearson R2 ≥ 0.5 and p-adjusted value ≤ 0.05, identifying 41 “positive regulators” (Figure 5F).

In conclusion, analysis of RNA and ATAC data of SFs in healthy and *hTNFtg* mice at single cell resolution, led to the identification of 9 sub-populations with distinct functions. Inspection of marker genes, enriched functional terms, marker peaks, enriched motifs and regulatory networks from both analyses supported the categorization of the identified clusters in 3 broad groups, namely sublining, intermediate and lining. In the sublining group, clusters showcase gene expression and accessibility patterns related to homeostatic properties, while clusters belonging to intermediate and lining groups differ at many aspects from the previously described category, and exhibit properties related to proliferation, inflammation and destruction of the joint. With the current use case, the complementarity of the two assays was showcased. To this end, results from the ATAC analysis were utilized in order to corroborate findings from the RNA such as cluster-specific TFs, marker genes and trajectories.

## DISCUSSION

SCALA is a comprehensive bioinformatics pipeline offered both as a web-server and a stand-alone application. It performs end-to-end scNGS analysis, by using the current best practices of the field. It currently enables the analysis of scRNA-seq and scATAC-seq datasets, which comprise the vast majority of the available scNGS data in the literature. Using state-of-the-art analysis modules and visualization, SCALA aids researchers to decipher complex biological mechanisms in an easy and convenient way. It handles both raw and pre-processed datasets, thus giving the users the flexibility to perform their analysis from-scratch, or to visualize and reanalyze already processed data. For comparison, we report a catalog of similar tools (e.g., pagoda2, SingleCAnalyzer (55), Bingle-seq (56), iCellR (57), cerebro (29), Is-CellR (58), SeuratWizard (31), ICARUS (59), SC1 (60), alona (61), WASP (62), CHIPSTER (63), Asc-Seurat (64), GenePattern (65), PIVOT (66) and we highlight their pros and cons along with their complementarity to SCALA (Supplementary Table 1). To this end, it is worth mentioning that to our knowledge, only icellR (67) offers scATAC seq analysis whereas only six applications are available as web server applications. In addition, SCALA is the only tool that offers L-R and GRN analysis modes. Conclusively, we expect that due to its simplicity and its pipeline integration, SCALA will become a reference application for biomedical scientists who wish to analyze and explore their data in an interactive way.

---

**Supplementary Table 1:**
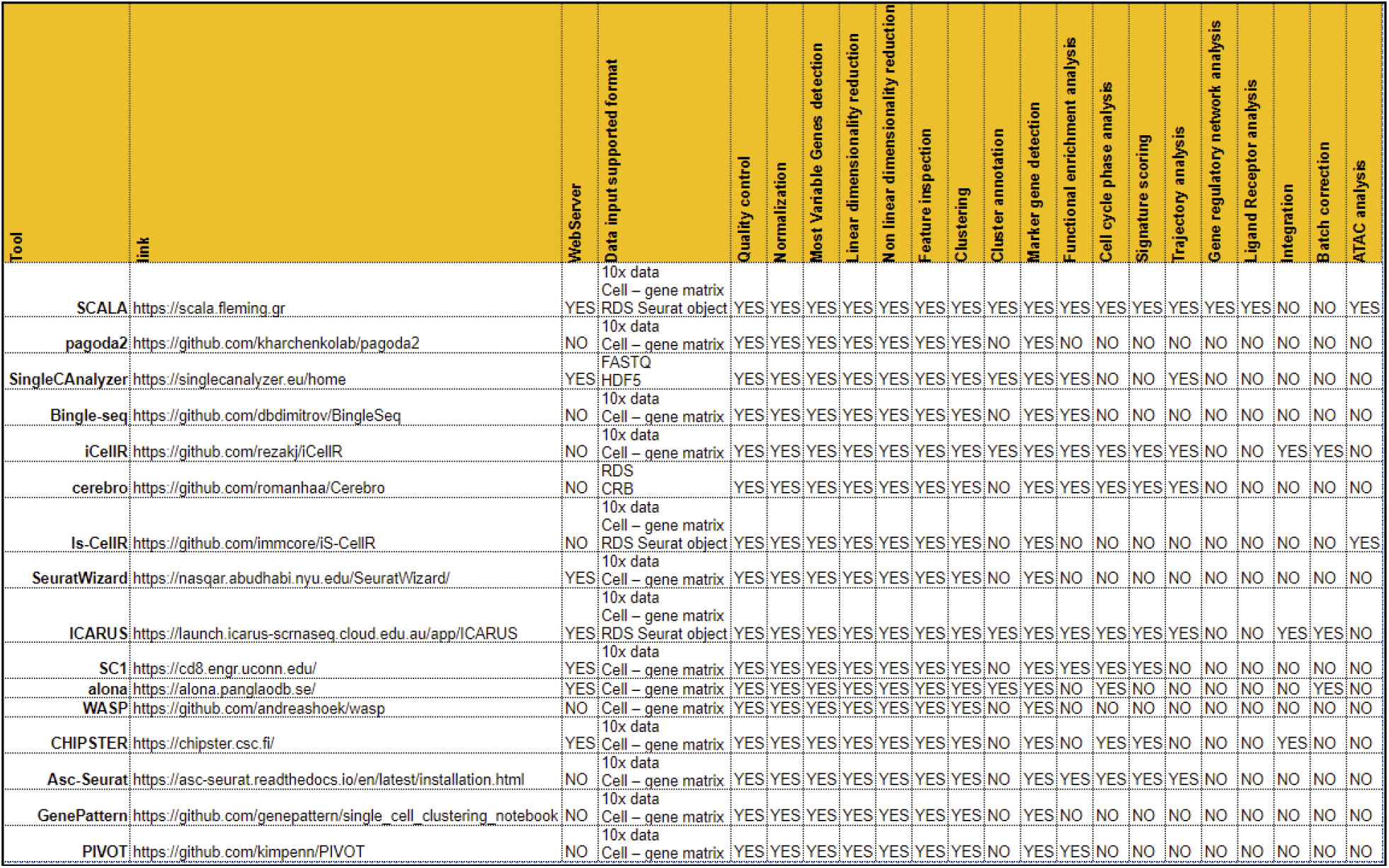
Comparison with similar tools.

## DATA AVAILABILITY

SCALA is an open source application and its code can be downloaded for local installation from the GitHub repository: https://github.com/PavlopoulosLab/SCALA. The service is also available online as a web application at http://scala.pavlopouloslab.info or https://scala.fleming.gr.

## AUTHOR CONTRIBUTIONS

CT and DK wrote most of the code. EK organized the GUI and worked on data visualization, FB implemented the functional enrichment analysis and worked on the back-end server and pipeline setup. DK implemented the pipeline related to scATAC-seq analysis. CT implemented the pipeline related to scRNA-seq analysis. CT, DK and GK provided the datasets for the case studies. DK, GK and GAP conceived the idea and supervised the project. All of the authors wrote parts of the manuscript.

